# Turnip crinkle virus targets host ATG8 proteins to attenuate antiviral autophagy

**DOI:** 10.1101/2021.03.28.437395

**Authors:** Aayushi Shukla, Gesa Hoffmann, Daniel Hofius, Anders Hafrén

## Abstract

Autophagy has emerged as a central player in plant virus disease and resistance. In this study we have addressed the potential roles of autophagy in Turnip crinkle virus (TCV) infection. We found that autophagy attenuates disease severity and contributes to resistance against TCV by limiting virus accumulation. These autophagy-dependent disease phenotypes intensify further when combined with defects in RNA silencing, suggesting that these two major defence pathways are largely uncoupled in TCV disease. Intriguingly, as a counterdefence, TCV employs the viral silencing suppressor protein P38 to suppress antiviral autophagy, likely by directly sequestering ATG8 proteins. This strategy appears to be novel for plant viruses, yet resembles mechanisms described for other pathogen classes. Together, these results broaden our understanding of autophagy in plant virus disease, and strengthens our view of virus-specific adaptation to the autophagy pathway.

## INTRODUCTION

Autophagy is a conserved intra-cellular degradation pathway that is essential for cells to recycle their macromolecules and organelles. While autophagy maintains at basal levels cellular homeostasis, tissue development and organelle quality control (Kraft and Martens, 2012), it can be substantially induced to facilitate adaptation to stress conditions like nutrient starvation, age-related senescence and pathogen attack (Boya et al., 2013; Klionsky and Codogno, 2013). Cytoplasmic targets of autophagy are engulfed in specialized double-membrane vesicles, called autophagosomes, that fuse with lysosomal or vacuolar compartments for lytic breakdown. The complex process of autophagosome biogenesis and turnover is controlled by more than 30 AuTophaGy related (*ATG*) genes that encode the core autophagy machinery conserved from yeast to mammals (Mizushima et al., 2011). There is ample evidence supporting specificity in governing the choice of cargo, known as selective autophagy, which requires efficient recognition of the specific cellular targets by autophagy receptors (Stephani and Dagdas, 2020; Zaffagnini and Martens, 2016). Selective autophagy can target cargo such as aggregated proteins, damaged or over-abundant organelles, and invading pathogens.

In recent years, autophagy has been implicated plant-pathogen interactions including diseases caused by viruses, bacteria, fungi and oomycetes (Hofius et al., 2017; Kushwaha et al., 2019; Leary et al., 2019). In plant-virus interactions, a plethora of examples have demonstrated that autophagy can have both anti-viral and pro-viral activities (Fu et al., 2018; Hafren et al., 2017; Hafren et al., 2018; Haxim et al., 2017; Ismayil et al., 2020; Li et al., 2018; Li et al., 2020b; Shukla et al., 2020; Yang et al., 2018). For instance, autophagy targets viral components and contributes to plant resistance via the cargo receptor NEIGHBOUR OF BRCA1 (NBR1) that mediates xenophagic degradation of *Cauliflower mosaic virus* (CaMV) viral particles (Hafren et al., 2017). Interestingly, autophagy can both support and limit virus infections of *Turnip mosaic virus* (TuMV) with divergent roles of NBR1 (Hafren et al., 2018; Li et al., 2018; Li et al., 2020a), clearly outlining autophagy as a central process with multifunctional, balancing capacities in virus disease. Adding to this complexity, viruses have acquired mechanisms to modulate autophagy in multiple ways including induction, suppression and subversion of its functions (Kushwaha et al, 2019). For example, the γb protein of *Barley stripe mosaic virus* (BSMV) directly binds to ATG7 and thus disrupts the ATG7-ATG8 interaction to suppress autophagy (Yang et al., 2018), while CaMV uses the viral P6 protein to down-regulate autophagy via dampening of salicylic acid responses (Zvereva et al., 2016).

RNA silencing is a major anti-viral pathway that involves processing of double-stranded RNAs (dsRNA) by DICER-LIKE (DCLs) proteins into small interfering RNAs (siRNAs). siRNAs are then integrated via binding to ARGONAUTE proteins (AGOs) into the RNA-induced silencing complex (RISC) to guide cleavage of viral RNAs. Viral Silencing Suppressors (VSRs) have been evolved by viruses to establish successful infections and are able to obstruct every step of the RNA silencing pathway (Csorba et al., 2015; Shukla et al., 2019). A crosstalk between autophagy and RNA silencing in viral infections has emerged in recent years. The polerovirus VSR P0 targets the RISC component AGO1 and triggers its ubiquitination and autophagic degradation (Derrien et al., 2012). On the contrary, autophagy favors plant defense and targets VSRs such as the potyviral helper-component proteinase (HCpro), cucumoviral 2b, and geminiviral bC1 for degradation (Hafren et al., 2018; Haxim et al., 2017; Ismayil et al., 2020; Nakahara et al., 2012; Shukla et al., 2020). At the same time, VSRs are commonly strong virulence factors and their autophagic degradation especially at later stages of infection should contribute to disease attenuation in favor of both the plant and virus.

*Turnip crinkle virus* (TCV) belongs to the genus *Carmovirus* of the family *Tombusviridae*. TCV is a small icosahedral positive-strand RNA virus that encodes five proteins. The capsid protein (CP) of TCV is a 38 kDa multifunctional protein (P38) required for virus assembly and suppression of RNA silencing (Choi et al., 2004). P38 also has a functional role the in long-distance movement of the virus (Cao et al., 2010). In this paper, we investigated the effect of autophagy on viral accumulation and development of disease in response to TCV infection in *Arabidopsis thaliana*, a compatible and natural host of TCV. We show that autophagy contributes to TCV resistance and suppression of virus-induced disease independent of the RNA silencing pathway. Intriguingly, we found that the P38 directly interacts with the core autophagy protein ATG8 to potentially suppress autophagy, outlining autophagy as a battle ground of TCV resistance.

## RESULTS

### Autophagy is activated and functional during TCV infection

We first monitored if autophagy is induced and altered during TCV infection. To this end, we used the stably expressing Arabidopsis lines GFP-ATG8a and NBR1-GFP, which are markers for autophagosomes and NBR1-dependent selective autophagy, respectively. Distribution of both markers was altered in TCV infected leaf tissue that showed a significantly higher number of GFP-ATG8a and NBR1-GFP puncta in systemically infected leaves (Fig.1 A and C). We used the commonly applied concanamycin A (ConA) treatment to inhibit the vacuolar ATPase and thereby stabilize autophagic bodies in the vacuole (Kataoka et al., 1996). Upon ConA treatment, GFP-ATG8a puncta were further increased in systemically infected leaf tissue compared to the DMSO control (Fig. 1 B and D), indicating that TCV triggers the formation of autophagosomes that are efficiently delivered for vacuolar degradation. GFP-ATG8 is cleaved after lysis of the autophagosome and the contents are exposed to vacuolar hydrolases. An intact GFP moiety is released as a result of the proteolysis of GFP-ATG8, which accumulates in the vacuole as it is relatively degradation-resistant. Therefore, the level of free GFP level correlates with autophagic flux (Nair et al., 2011). In accordance with the microscopy analysis, the level of free GFP was also higher in TCV infected leaf tissue as compared to the non-infected controls (Fig 1 E), together supporting enhanced autophagic flux during TCV infection.

**Figure 1.**
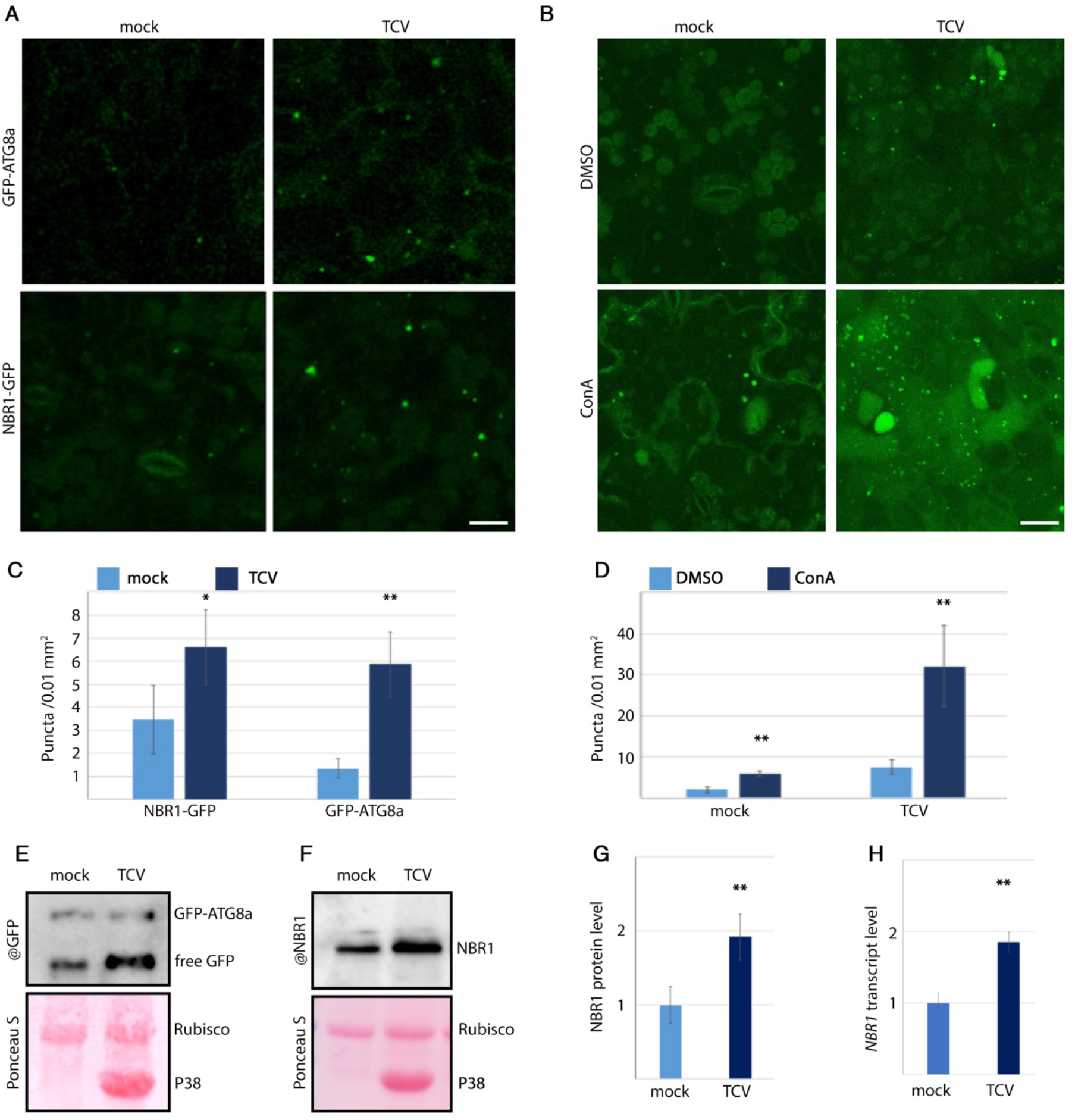
Autophagy induction after TCV infection. A. Representative images of GFP-ATG8a and NBR1-GFP marker in uninfected control and TCV infected plants. B. Representative images of GFP-ATG8a marker in uninfected control and TCV infected plants after DMSO or ConA treatment. Images are projected as confocal Z-stacks. Scale bars = 10 *μ*m in (A and B). C and D. GFP-ATG8a puncta counted from similar images as in A and B using ImageJ (*n* = 9). E. Anti-GFP western blot analysis to estimate free GFP levels in uninfected control and TCV-infected GFP-ATG8a plants. Ponceau S staining verified loading control. F. Anti-NBR1 western blot analysis for detection of NBR1 levels in uninfected control and TCV infected Col-0 plants. Ponceau S verified loading control. G. Protein levels of NBR1 in uninfected control and TCV infected Col-0 as compared to Rubisco, analysed using ImageJ after immunodetection from four replicate samples as in (F). H. Transcript levels of *NBR1* in control and TCV infected Col-0 normalized by *PP2a* as determined by RT-qPCR (*n* = 4). Statistical significance (\**P* < 0.05; \*\**P* < 0.005) calculated by analysis of variance and pairwise comparison by least significant difference (LSD) test.

The selective autophagy receptor NBR1 is recruited by the developing autophagosomes to deliver cargo through interaction with ATG8 proteins. Due to its degradation along with autophagosomes, NBR1 protein accumulation serves as another indicator of autophagy activity and flux (Klionsky et al., 2021; Svenning et al., 2011). We found that NBR1 accumulated to higher levels during TCV infection compared to the non-infected control plants (Fig. 1 F and G). Because *NBR1* transcript levels were elevated to a similar extent during infection (Fig. 1 H), this finding further supported our notion that autophagic flux is functional during TCV infection in Arabidopsis. Altogether, these assays showed that autophagy is activated and functional during TCV infection.

### Autophagy suppresses disease severity and promotes resistance against TCV infection

All major plant defence strategies against pathogens impact the fitness of the plant, which is influenced by severity of disease caused by the pathogen. Biomass loss can be a measure of disease severity or virulence upon virus infection. Previously, we showed that the core autophagy genes *ATG5* and *ATG7* are important for reducing disease severity in CaMV, TuMV and CMV infected plants, while the selective autophagy cargo receptor *NBR1* had no significant effect (Hafren et al., 2017; Hafren et al., 2018; Shukla et al., 2020). To address whether functional autophagy affects TCV disease development in a similar manner, we monitored infection development in loss of function mutants *atg5* and *nbr1* compared to the Col-0 wild-type (WT). At 28 days post inoculation (DPI), Col-0 WT and *nbr1* plants showed typical TCV symptoms including stunting, leaf crinkle and chlorosis, and a similar reduction in biomass, while *atg5* plants were noticeably more affected by developing senescence symptoms and more pronounced biomass loss (Fig. 2 A and B). This observation is in accordance with the previous findings during CaMV, TuMV and CMV infection, thus strengthening the general importance of autophagy in plant tolerance to viruses.

**Figure 2.**
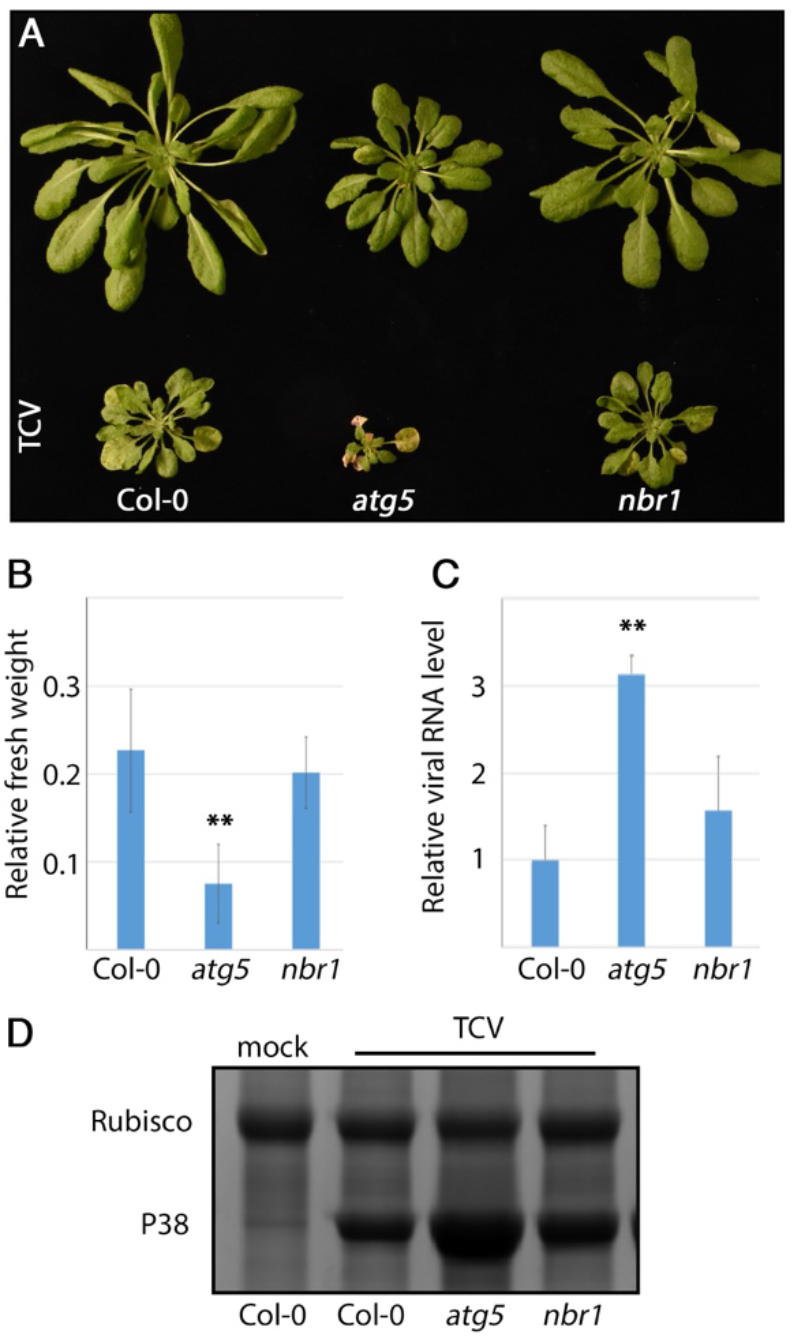
Autophagy promotes plant resistance against TCV infection. A. Representative images of control (upper row) and infected (lower row) Col-0, *atg5, nbr1* at 28 DPI. B. The relative fresh weight of TCV infected plants to uninfected controls at 28 DPI (*n = 10*). C. Relative TCV RNA levels determined by RT-qPCR at 28 DPI in different genotypes (*n* = 4). Statistical significance (\*\**P* < 0.005) calculated by analysis of variance and pairwise comparison by least significant difference (LSD) test. D. Coomassie Brilliant Blue (CBB) staining was used to assess the accumulation of TCV coat protein P38 in denoted genotypes at 28 DPI. Accumulation of the Rubisco large subunit served as loading control.

To assess whether autophagy plays a role in resistance against TCV, we compared viral RNA accumulation levels in *atg5, nbr1*, and Col-0 WT plants (Fig. 2C). Intriguingly, TCV RNA accumulated to significantly higher levels in *atg5* than in WT and *nbr1*. Likewise, the highly abundant capsid protein P38 that is easily detected in total protein stains, was clearly more abundant in *atg5* than the others genotypes (Fig. 2D). Together, these results revealed that autophagy functions in resistance and attenuates disease symptoms during TCV infection.

### Autophagy and RNA silencing contribute independently to TCV resistance and disease suppression

RNA silencing is a major defence mechanism of plants that viruses frequently counteract by evolving RNA silencing suppressors (Csorba et al., 2015). TCV P38 is a silencing suppressor that obstructs the dicer-like proteins (DCL2/DCL4) in the host plant to overcome RNA silencing (Thomas et al., 2003) Because P38 accumulated to higher levels in *atg5*, one possibility is that the silencing suppression capacity is increased in *atg5* causing compromised resistance. Therefore, we investigated the possible epistasis between RNA silencing and autophagy by comparing the higher-order mutants *atg7 dcl2 dcl4* and *atg5 ago1* to the *atg5, atg7, ago1* and *dcl2 dcl4* background lines. The triple mutant *atg7dcl2dcl4* showed an extreme growth reduction, severe chlorosis and early senescence phenotype during TCV infection by 28 DPI, being evidently additive between *atg7* and *dcl2 dcl4* (Fig. 3A and B). While viral RNA levels were elevated in both *atg7* and *dcl2 dcl4*, we omitted the triple mutant owing to its severe disease (Fig. 3E). However, we speculate that the triple mutant accumulates viral RNA additively, as revealed for P38 protein levels in a total protein stain (Fig. 3F).

**Figure 3.**
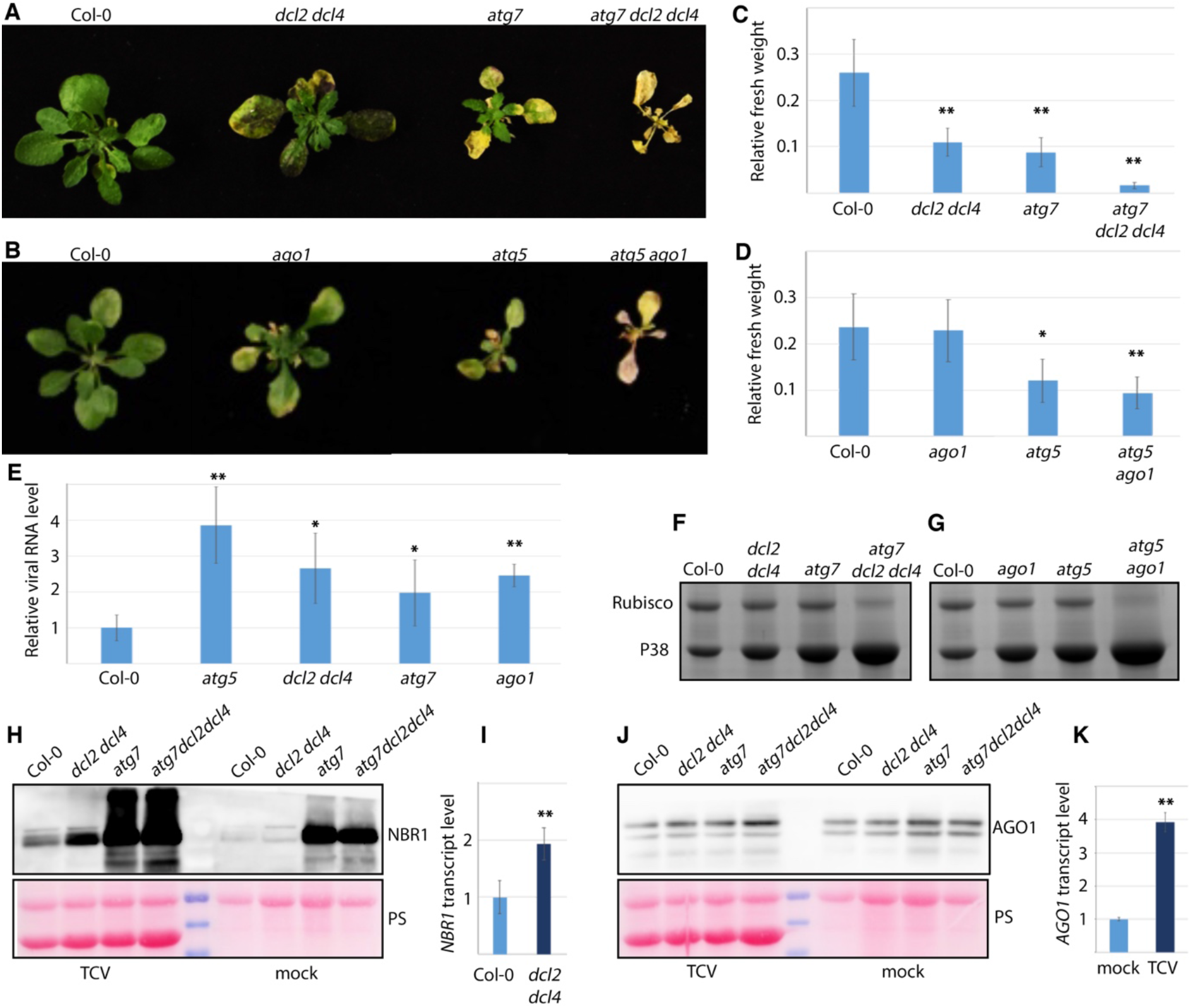
Additive effect of autophagy and RNA silencing on TCV P38 protein accumulation. A. and B. Representative images of infected genotypes (Col-0, *ago1, atg5, atg7, atg5ago1, dcl2dcl4, atg7dcl2dcl4*) at 21 DPI. C. and D. The relative fresh weight of TCV infected plants to uninfected controls at 21 DPI (*n* = 10). E. Relative TCV RNA levels determined by RT-qPCR at 21 DPI in different genotypes (*n* = 4). F. and G. Coomassie Brilliant Blue (CBB) staining was used to assess the accumulation of TCV coat protein P38 in different genotypes at 21 DPI. Accumulation of the Rubisco large subunit served as a loading control. H. Anti-NBR1 western blot analysis for detection of NBR1 levels in uninfected control and TCV infected Col-0, *dcl2dcl4, atg7, atg7dcl2dcl4* genotypes at 13 DPI. Ponceau S (PS) verified loading control. I. Transcript levels for *NBR1* in TCV infected Col-0 and *dcl2dcl4* at 13 DPI, determined by RT-qPCR (*n* = 4) J. Anti-AGO1 western blot analysis for detection of AGO1 levels in uninfected control and TCV infected Col-0, *dcl2dcl4, atg7, atg7dcl2dcl4* genotypes at 13 DPI. Ponceau S (PS) verified loading control. K. Transcript levels for *AGO1* in uninfected control and TCV infected Col-0 and at 13 DPI, determined by RT-qPCR (*n* = 4). Statistical significance (\**P* < 0.05; \*\**P* < 0.005) calculated by analysis of variance and pairwise comparison by least significant difference (LSD) test.

In addition to suppressing DCL2 and DCL4, P38 is also known to bind the slicer AGO1 as part of counteracting the RNA silencing pathway (Azevedo et al., 2010). We observed a similar lack of epistasis between *atg5* and *ago1* as seen for the dicer mutant *dcl2 dcl4* when infected with TCV. At 21 DPI, the biomass loss was significantly higher in the *atg5* and *ago1* single mutants, and the *atg5 ago1* double mutant further showed an additive disease phenotype including escalated senescence (Fig. 3D). Similar to the *atg7 dcl2 dcl4* triple mutant, we could not explore TCV RNA accumulation in *atg5 ago1* owing to its early senescence phenotype after TCV infection (Fig. 3C). However, P38 protein accumulation was still detectable and showed additivity between the already higher levels present in both *atg5* and *ago1* single mutants (Fig. 3 G). The additive disease phenotype in *atg7 dcl2 dcl4* and *atg5 ago1* was also revealed by the strongly reduced levels of the large Rubisco subunit in the protein-stained gels (Fig. 3F and G).

These results support the view that autophagy and RNA silencing act independently to suppress P38 accumulation. Nonetheless, we still wanted to rule out that autophagy was defective in the *dcl2 dcl4* background and that the autophagic degradation of AGO1 (Derrien et al., 2012) was activated by TCV, thereby mediating P38 degradation through direct AGO1 interaction (Azevedo et al., 2010; Jin and Zhu, 2010). We analysed NBR1 protein levels in Col-0 WT and the mutants *dcl2 dcl4*, *atg7* and *atg7dcl2dcl4* in non-infected and TCV-infected conditions (Fig. 3H). Indeed, we could observe higher levels of NBR1 in *dcl2 dcl4* after TCV infection as compared to the corresponding non-infected controls, and interestingly, NBR1 transcript levels were likewise increased by several fold (Fig. 3I). This finding suggested that autophagy is rather induced than impaired in the *dcl2 dcl4* background. To evaluate whether TCV activated the autophagic degradation of AGO1, we determined AGO1 proteins levels at 13 DPI in TCV infected plants in comparison to non-infected controls (Fig. 3J). As we did not observe over-accumulation of AGO1 in the *atg7* background, the autophagic degradation of P38 through AGO1 interaction appears unlikely. Notably, despite comparable AGO1 levels between uninfected control and infected plants, *AGO1* transcript levels were increased by 4-fold in TCV infected compared to non-infected plants (Fig 3K). These results indicated the autophagic degradation of P38 is AGO1 independent.

### P38 interacts with ATG8 proteins to suppress autophagy

Next, we aimed to dissect the autophagy-based resistance mechanisms that restricts viral RNA and capsid protein P38 accumulation. We, therefore, checked virus accumulation in the mutant lines of several known selective autophagy receptors. These included the ER-associated receptor mutant *ati1 ati2* (Sjøgaard et al., 2019); the quadrupole *pux-q* mutant (Marshall et al., 2019) and the proteaphagy receptor *rpn10* (Marshall et al., 2015) in addition to *nbr1*. At 21 DPI, viral RNA accumulation did not differ in any of the receptor mutant lines (Fig. 4A), which led us to assess whether P38 could be directly interacting with ATG8 proteins. Interestingly, we found that all four tested RFP-tagged TCV proteins; P8, P9, P28 and P38, co-localized with both NBR1-GFP and GFP-ATG8a upon co-expression in *Nicotiana benthamiana* (Fig. S1). This finding may suggest a general role of autophagy in TCV protein turnover; however, the level of co-localization differed for the viral proteins and was most evident for P38 in combination with the ATG8 protein. Therefore, we established transgenic lines expressing P38-RFP in the GFP-ATG8a background. Indeed, these lines verified co-localization of GFP-ATG8a and P38-RFP in cytoplasmic aggregates of variable size (Fig. 4B). More excitingly, when this line was treated with ConA we did not detect P38-RFP in the vacuole (Fig.4B) and compared to the GFP-ATG8a control, the number of autophagic puncta was clearly reduced by P38-RFP (Fig. 4C). Together with the notion that the number of P38-RFP aggregates co-localized with GFP-ATGG8a were not increased by ConA treatment (Fig.4C), it appeared rather that P38 could suppress autophagy by sequestering ATG8 directly or via other ATG8-interacting components into aggregates.

**Figure 4.**
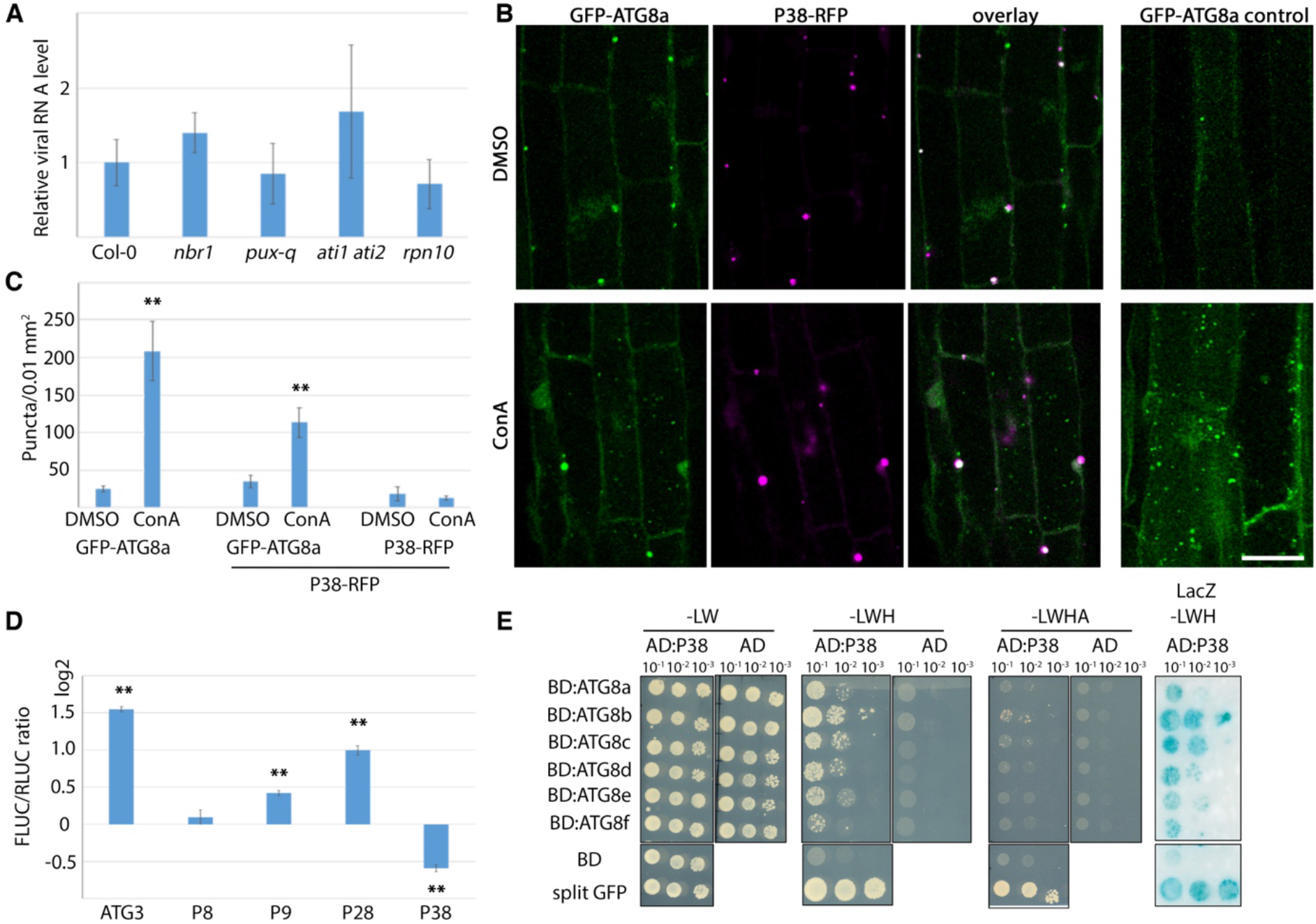
P38 interacts with ATG8 to suppress autophagy. A. Relative TCV RNA levels determined by RT-qPCR at 21 DPI in selected known autophagy receptor mutants (*n* = 4). B. Representative single-plain images of colocalization of P38-RFP (Magenta) with GFP-ATG8a (Green) in the roots of 12 day old seedlings of Arabidopsis transgenic line P38-tagRFP and GFP-ATG8a after DMSO or ConA treatment. GFP-ATG8a seedlings were used as control. Scale bar = 20 *μ*m. C. GFP-ATG8a and P38-RFP puncta counted from similar images as in B using ImageJ (*n* = 9). D. RLUC-ATG8a and the internal control FLUC were co-expressed with TCV proteins in *N. benthamiana*. ATG3 and GUS were used as control. Values represent the mean ratio of RLUC-ATG8a and FLUC activities, measured 72 h post infiltration in the dual-luciferase system. Statistical significance (\**P* < 0.05; \*\**P* <0.005) calculated by analysis of variance and pairwise comparison by least significant difference (LSD) test. E. Y2H assay for interaction of P38 fused to activation domain (AD) and different ATG8 isoforms fused to binding domain (BD). Split GFP (BD: 1-10-GFP / AD: 11-GFP) was used as a positive control. -LWHA (-leu/trp/his/ade).

This observation prompted us to analyse effects of individual TCV proteins on autophagy levels using a recently developed quantitative assay in *N. benthamiana* leaves, that is based on the transient expression of ATG8a fused to *Renilla* luciferase (RLUC) together with firefly luciferase (FLUC) as internal control (Leong et al., 2021; Ustun et al., 2018). Co-expression of the viral protein P38 significantly increased RLUC-ATG8a accumulation compared to the FLUC control, suggesting autophagy inhibition (Fig.4D). The opposite was observed for P28, which significantly decreased RLUC-ATG8a accumulation in relation to FLUC. The other proteins behaved more similar to the GUS control. Over-expression of ATG3 was previously shown to induce autophagy in *N. benthamiana* (Han et al., 2015) and indeed this control resulted in a 4-fold increase of FLUC to RLUC-ATG8a ratio supporting functionality of the assay (Fig. 4D). Together, this result strengthened P38 as a suppressor of autophagy, and identified P28 as a potential autophagy inducer.

All our observations with P38 could be explained by a direct interaction with ATG8s. To assess this possibility, we used the yeast-2-hybrid (Y2H) system. We fused P38 with the GAL-4 activation domain (AD) and six ATG8 isoforms (ATG8a-ATG8f) with the GAL-4 binding domain (BD). Intriguingly, P38 showed positive interaction with all the tested ATG8 isoforms as revealed by growth on -leu/trp/his medium and the LacZ assay (Fig. 4E). Taken together with the quantitative assays and that the co-localized aggregates of P38 and ATG8 do not appear to be prominent autophagy targets, we propose that P38 directly interacts with several ATG8 isoforms to potentially suppress autophagy.

## DISCUSSION

Similar to other plant-pathogen interactions, there is a continuous molecular arms race between resistance and virulence during virus infection. Every defence response of plants eventually evokes a counter-defence mechanism by the virus, putting both under different selective pressures. Many studies have established that autophagy has evolved as a frontrunner in plant virus defence and tolerance, with a broad-range of mechanisms deduced collectively for Caulimoviruses (Hafren et al., 2017), Potyviruses (Hafren et al., 2018; Li et al., 2018), Hordeiviruses (Yang et al., 2018), Tenuiviruses (Fu et al., 2018), Geminiviruses (Haxim et al., 2017; Ismayil et al., 2020) and Cucumoviruses (Shukla et al., 2020). Given the important role of autophagy in maintaining plant homeostasis, cellular quality control and stress adaptation, it is understandable that autophagy is a key contributor in virus defence. Autophagy can mediate resistance by selectively degrading viral components, but several studies have also indicated that distinct viruses have evolved mechanisms to counteract, modulate and exploit autophagic processes to promote virulence. Hence, it became evident that both anti- and proviral functions of autophagy operate in parallel during virus infections, more extensively revealed in animals but ample evidence is also building up in plants (Kushwaha et al., 2019; Levine and Kroemer, 2019)

In this study we found that autophagy restricts TCV accumulation and that P38 associates closely with ATG8 proteins. Because P38 is the RNA silencing suppressor of TCV (Iki et al., 2017; Thomas et al., 2003), one possibility was that autophagy degrades P38 to potentiate this antiviral defence system, thereby enhancing resistance. However, higher-order mutants of autophagy and RNA silencing suggested lack of epistasis of these pathways in TCV resistance by showing additivity in P38 accumulation. This observation implied that resistance provided by autophagy and RNA silencing are uncoupled and independently suppress TCV accumulation, similar to TuMV and opposite to CMV (Hafren et al., 2018; Shukla et al., 2020). Xenophagy is a type of selective autophagy that is utilized against invading pathogens by their entire elimination (Mao and Klionsky, 2017). The targeting and degradation of CaMV particles via the cargo receptor NBR1 represents one of the primary examples of xenophagy in plants (Hafren et al., 2017), showing striking similarities to particle degradation of several animal viruses through the corresponding receptor p62 (Berryman et al., 2012; Judith et al., 2013; Orvedahl et al., 2010; Orvedahl et al., 2011; Shelly et al., 2009). Considering the interaction between ATG8s and the capsid protein P38 combined with the high accumulation of P38 and viral RNA in core autophagy mutants, viral particle xenophagy is a possible explanation. However, this requires further evidence as it is still equally possible that autophagy limits virus replication and thus has a more indirect impact on virus particle accumulation.

Despite the clear co-localization of P38 and ATG8 in cytoplasmic aggregates, we could not detect the clearance of these structures by autophagic degradation in P38 transgenic lines, questioning efficient P38 xenophagy at least outside of the infection context. Instead, we uncovered the intriguing capacity of P38 to suppress autophagy, which we now hypothesize is mediated via direct sequestration of ATG8s. ATG8 (LC3 in animals) is a ubiquitin-like protein that functions as a conjugate with the phospholipid PE (ATG8–PE) during autophagy. ATG8 contributes to ATG1 recruitment, is essential for autophagosome membrane formation and involved in cargo selectivity through its interaction with cargo receptors (Farre and Subramani, 2016). Recently, our understanding of ATG8-dependent recognition of receptor proteins and cargo has hugely expanded (Klionsky and Schulman, 2014). ATG8 is the most diversified protein among the core ATG proteins (Bu et al., 2020) and its central importance for the autophagy process exposes ATG8 as a potential effector target. Indeed, different effector proteins from bacteria, fungi, oomycetes and nematodes can interact with ATG8s (Lal et al., 2020). Evidently, the interaction of a viral component with ATG8 may result in autophagic degradation to influence infection, as observed for the *Cotton leaf curl Multan virus* (CLCuMuV) virulence factor bC1, which is targeted for autophagic degradation through its direct interaction with ATG8f (Haxim et al., 2017). An alternative, and not mutually exclusive outcome could be that an interaction with ATG8 results in the suppression of autophagy and associated resistance to thrive infection. A similar mechanism was revealed for the unrelated plant and animal pathogen effectors HopF3 of *Pseudomonas syringae* (Lal et al., 2020), RavZ of *Legionellla pneumophila* (Choy et al., 2012), and PexRD54 of *Phytophthora infestans* (Dagdas et al., 2018). We propose that the interaction of P38 with several ATG8s has evolved to mediate suppression of autophagy. The manipulation of ATG8 to inhibit autophagy has not been previously shown for plant viruses, thereby identifying another mechanisms in the intricate interaction with their hosts. Excitingly, the auxillary replication protein P28 was a strong inducer of autophagy in our quantitative assay, which may well unintendedly trigger autophagy-based resistance that needs to be balanced by the suppressing activities of P38 to facilitate strong infection and virus accumulation. Our findings underscore the high complexity in the interaction of TCV with autophagic processes and further support the emerging view that viral proteins may both trigger and block autophagy for the benefit of infection.

## MATERIAL AND METHODS

### Plant material and growth conditions

Wild-type (WT) plant was *Arabidopsis thaliana* ecotype Columbia (Col-0). Mutants *atg5-1, nbr1, ago1-27, dcl2 dcl4, atg7 dcl2 dcl4*, and the GFP-ATG8a and NBR1-GFP transgenic line have been described previously (Hafren et al., 2018). The *ago1-27 atg5-1* double mutant was generated by crossing, as described in (Shukla et al., 2020). The transgenic line expressing P38 tag-RFP (pGWB660) was obtained by floral dip transformation of the GFP-ATG8a line. For infection experiments, Arabidopsis plants were grown on vermiculite soil under short-day conditions (8/16-h light/dark cycles) at light intensity of 150 μE/m^2^s and 21°C in a growth chamber. For transient expression assays, *N. benthamiana* plants were cultivated under long-day conditions (16/8-h light/dark cycles) in a growth chamber at 150 μE/m^2^s light intensity, 21°C temperature, and 70% relative humidity.

### DNA constructs

TCV viral proteins P8, P9, P28 and P38 were amplified using plasmid pTCV66, which contains the full-length wild type TCV (TCV-M strain) sequence located downstream of a T7 RNA polymerase promotor. The plasmid was used as template and cloned into pENTRY-DTopo and further recombined into gateway vector pGWB660 using LR Clonase II (Invitrogen Life Technologies). Expression constructs for GUS and GFP-ATG8a were described previously in (Hafren et al., 2018). All binary vectors were transformed into *Agrobacterium* C58C1 for transient expression in *N. benthamiana* or floral dip transformation of Arabidopsis (Clough and Bent, 1998).

### TCV infection and quantification

TCV-M transcripts were obtained using plasmid pTCV66, linearized with *Sma*I followed by in-vitro transcription using T7 RNA polymerase. *N. benthamiana* plants were then inoculated with the in-vitro transcripts. After 12 days, sap of the infected leaves was used to prepare inoculum using phosphate buffer (0.5 M Na_2_HPO_4_, 1 M NaH_2_PO_4_ pH 7.0) with 0.2 % diethyldithiocarbamic acid (DIECA) as a reducing agent (Shukla et al., 2018). The first two true leaves of 3-week-old Arabidopsis plants [stages 1.04–1.05 as in (Boyes et al., 2001)] were sprinkled with 0.2 % carborundum and mechanically inoculated with 5 μl of the inoculum per plant. Plants were sampled in biological replicates, each containing 3 individual plants from which inoculated leaves were removed. Subsequent leaves from the uninoculated plants were also removed before harvesting the control. For TCV RNA or plant transcript quantitation, total RNA was isolated using the method as described in (Onate-Sanchez and Vicente-Carbajosa, 2008). First-strand cDNA was obtained using Maxima First Strand cDNA Synthesis Kit (Thermo Fisher Scientific). Quantitative RT-PCR analysis (qPCR) was performed with Maxima SYBR Green/Fluorescein qPCR Master Mix (Thermo Fisher Scientific) using the CFX Connect™ Real-Time PCR detection system (BIO-RAD) with gene-specific primers listed in Supplemental Table 1. Normalization was done using *PP2A* (*AT1G69960*).

### Confocal microscopy and treatments

Live cell images were acquired from roots cells or abaxial leaf epidermal cells using Zeiss LSM 800 microscope and processed with ZEN Blue software. Excitation/detection parameters for GFP was 488 nm/490-552 nm. Treatment with 0.5 μM concanamycin A (Santa Cruz Biotechnology) or DMSO was carried out in liquid ^1^/2 MS for 10 h before confocal analysis. Quantitation of GFP-ATG8a labelled puncta was done using Image J (version 2.1.0/1.53c). For the quantification of ATG8a labelled puncta, images were stacked using ‘Z-projection’ followed by ‘Gaussian blur’ to negate the background, and then puncta were counted under ‘Find maxima’ with a set threshold.

### Dual Luciferase Assay

The dual luciferase reporter assay was performed according to the manufacturer’s instructions (Dual-Luciferase Reporter Assay System; Promega) with slight modifications. Briefly, four leaf discs were homogenized in 200 μL lysis buffer and cleared by centrifugation. For detection and measurement of the Firefly luciferase activity, 35 μL of the luciferase assay reagent was added to 5 μL of plant extracts. To measure Renilla luciferase activity, 35 μL of the Stop and Glo reagent was added to the mixture. The measurement was performed in an Omega Fluostar plate reader.

### Immunoblot analysis

Proteins were extracted in 100 mM Tris pH 7.5 with 2% SDS, kept at 99°C for 5 min in Laemmli sample buffer and cleared by centrifugation. The protein extracts were then separated by SDS-PAGE, transferred to polyvinylidene difluoride (PVDF) membranes (Amersham, GE Healthcare), blocked with 5% skimmed milk in PBS, and incubated with primary antibodies anti-NBR1 (Svenning et al., 2011), anti-AGO1 (Agrisera; AS09 527) and anti-GFP (Santa Cruz Biotechnology; sc-9996) using 1:3000 dilution in PBS 0.1% Tween-20, and secondary horseradish peroxidase-conjugated antibodies 1:10 000 in PBS 0.1% Tween-20 (Amersham, GE Healthcare). The immunoreaction was developed using the ECL Prime kit (Amersham, GE Healthcare) and detected in a LAS-3000 Luminescent Image Analyzer (Fujifilm, Fuji Photo Film). For staining, SDS-PAGE gel was stained using Coomassie Stain solution (Coomassie Blue, Methanol, Acetic Acid) for 1 h on shaking at room temperature and destained using Destaining solution (Methanol, Acetic Acid) overnight shaking at room temperature.

### Yeast two-hybrid

Y2H techniques were performed according to the Yeast Protocols Handbook (Clontech). The gateway-compatible Y2H vectors PGBKT7 and PGADT7 were used to generate fusions of the ATG8 isoforms and TCV P38 respectively. Yeast strain AH109 was co-transformed with the respective plasmid combinations, including empty vector controls, followed by selection on solid double dropout (-Trp/Leu), triple dropout (-Trp/Leu/Ade) and quadruple dropout (-Trp/Leu/Ade/His) SD medium for 5 days at 28 °C. The interaction was analyzed by growth on quadruple dropout followed by lacZ assays.

### Data analysis and presentation

Data are presented as mean ± standard deviation (SD) and statistical significance was analyzed by ANOVA test followed by posthoc least significant difference (LSD) test. The obtained *p*-values <0.05 denoted * and *p*-values <0.01 denoted **. The number of replicates is given in the respective figure legends (*n*). All analysis has been performed using SPSS software v 23 (IBM Corp., USA).

## Acknowledgements

We gratefully acknowledge Anne Simon (University of Maryland, US) for providing the TCV plasmid pTCV66 strain M, Denise Seitner (Gregor Mendel Institute, Austria) for the positive control of yeast two hybrid, Richard Marshall and Richard Vierstra (Washington University, St. Louis, US) for the quadripole *pux-q* mutant, Hadas Peled-Zehavi (Weizmann Institute of Science, Rehovot, Israel) for the *ati1/2* double mutant, Jan Smalle (University of Kentucky, Lexington, US) for the *rpn10* mutant. Funding from FORMAS to A.H (grant number 2016-01044) and D.H. (grant number 2017-01596) is also gratefully acknowledged.

---

**Supplement figure 1.**
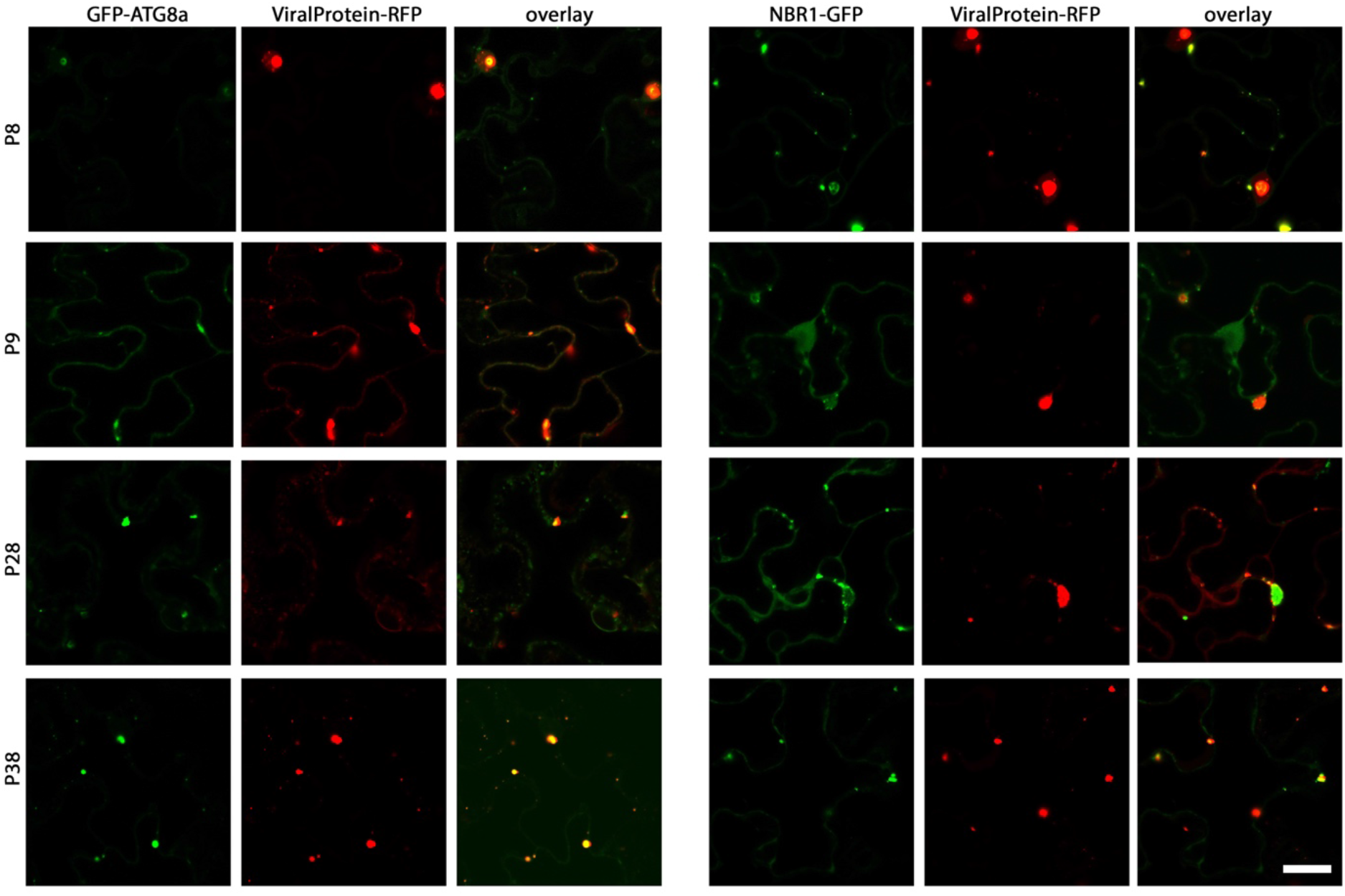
Co-expression of TCV proteins; P8-RFP, P9-RFP, P28-RFP and P38-RFP with GFP-ATG8a and NBR1-GFP in *N. benthamiana*. Images taken 48 h after infiltration. Scale bar = 20 *μ*m.

